# Intranasal delivery of lipid nanoparticle encapsulated SARS-CoV-2 and RSV-targeting siRNAs reduces lung infection

**DOI:** 10.1101/2022.07.25.501479

**Authors:** Aroon Supramaniam, Yaman Tayyar, Daniel. T. W. Clarke, Gabrielle Kelly, Kevin V. Morris, Nigel A. J. McMillan, Adi Idris

## Abstract

RNA interference (RNAi) is an emerging and promising therapy for a wide range of respiratory viral infections. This highly specific suppression can be achieved by the introduction of short-interfering RNA (siRNA) into mammalian systems, resulting in the effective reduction of viral load. Unfortunately, this has been hindered by the lack of a good delivery system, especially via the intranasal (IN) route. Here, we have developed an IN siRNA encapsulated lipid nanoparticle (LNP) *in vivo* delivery system that is highly efficient at targeting severe acute respiratory syndrome coronavirus 2 (SARS-CoV-2) and respiratory syncytial virus (RSV) in infected mouse lungs. Importantly, IN siRNA delivery without the aid of LNPs abolishes anti-SARS-CoV-2 activity *in vivo*. Our approach using LNPs as the delivery vehicle overcomes the significant barriers seen with IN delivery of siRNA therapeutics and is a significant advancement in our ability to delivery siRNAs. The studies presented here demonstrates an attractive alternate therapeutic delivery strategy for the treatment of both future and emerging respiratory viral diseases.

## Introduction

The novel coronavirus disease 2019 (COVID-19) caused by severe acute respiratory syndrome coronavirus 2 (SARS-CoV-2) has a mortality rate of ∼2.0% (1). Despite the deployment of several vaccines, SARS-CoV-2 variants continue to emerge, and there remains a need for a direct acting therapeutic. Similarly, direct acting antivirals against another emerging respiratory virus, respiratory syncytia virus (RSV) are also needed. RSV is the most common cause of bronchiolitis and pneumonia in children over 1 year of age. Research towards effective treatment and a vaccine against RSV has been ongoing for nearly five decades with little success. Current treatments for RSV including palivizumab (for high-risk children) and ribavirin (2) are limited and have poor efficacy resulting in a supportive rather than curative approach. Twenty months into the COVID-19 pandemic, we have only seen the first oral antiviral drug, molnupiravir, that shortens time to clearance of SARS-CoV-2 (3) and if administered early following infection, can reduce hospitalisation and death. With the advent of RNA biologics during the COVID-19 pandemic era and its rapid speed of development (e.g., Moderna mRNA-based COVID-19 vaccines (4)), there has been a strong impetus for the development of RNA-based therapeutics for a range of human diseases (5). RNA interference (RNAi) technology platform is rapidly regaining interest in the field of RNA biologics. As RNAi is modular and facile this approach could be applied to any virus of concern. RNAi works via the introduction of small interfering RNAs (siRNAs) that specifically target the protein-coding mRNAs via sequence complementarity, causing their subsequent degradation (6). RNAi is potentially cost effective, scalable and can be easily programmed to target any viral RNA in a matter of days. However, the greatest challenge is their delivery, which is particularly difficult when targeting the respiratory tract. We have recently demonstrated that siRNAs complexed in lipid nanoparticles (LNPs) designed to highly conserved regions of SARS-CoV-2 give potent viral lung repression *in vivo* when delivered intravenously (IV) (7). Given the limited clinical utility for delivering siRNA via IV, here we aim to develop an intranasal (IN) siRNA therapy targeting SARS-CoV-2 and RSV. This serves as a more clinically tractable way of ameliorating virus infection in the lungs. We hypothesize that RNAi can be deployed as a therapy to treat these viral infections using IN delivery approaches that target the respiratory epithelium suitable for outpatient use. Indeed, siRNA therapies against RSV have been attempted by Alnylam Pharmaceuticals with ALN-RSV01, a nasal spray formulation of a single naked siRNA directed against RSV nucleocapsid (N) protein gene (8). Similarly for SARS-CoV-2, there is precedence for this approach as IN delivery of naked (9) and dendrimer formulated (10) siRNAs are successful at ameliorating SARS-CoV-2 infectious load in lungs of infected animals. Here, we formulated previously designed siRNAs targeting SARS-CoV-2 (7) and an RSV-targeting siRNA in LNPs for IN delivery. The results presented here demonstrate that IN delivered LNP-complexed siRNAs effectively and significantly reduced SARS-CoV-2 and RSV load in lungs of infected mice.

## Materials and methods

### Cell culture

Vero E6 cells were maintained in complete media; DMEM (Gibco-Invitrogen, Waltham, MA) supplemented with 10% heat inactivated fetal bovine serum (FBS) (30 min at 56°C, Gibco-Invitrogen, Waltham, MA) and 1% of antibiotic/glutamine preparation (100 U/ml penicillin G, 100 U/ml streptomycin sulphate, and 2.9 mg/ml of L-glutamine) (Gibco-Invitrogen, Waltham, MA). A549 cells were maintained in complete media; DMEM/F12 (Gibco-Invitrogen, Waltham, MA) supplemented with 10% heat inactivated fetal bovine serum (FBS) (30 min at 56°C, Gibco-Invitrogen, Waltham, MA) and 1% of antibiotic/glutamine preparation (100 U/ml penicillin G, 100 U/ml streptomycin sulphate, and 2.9 mg/ml of L-glutamine) (Gibco-Invitrogen, Waltham, MA).

### Chemicals, lipids, and reagents

Cyclophosphamide was obtained from Sigma-Aldrich (St. Louis, MO). Dioleoyl trimethylammonium propane (DOTAP), dioleoylphosphatidylethanolamine (DOPE), 1,2-distearoyl-sn-glycero-3-phosphocholine (DSPC), Polyethylene Glycol (PEG)2000-C16 Ceramide conjugate, and cholesterol were purchased from Sigma-Aldrich (St Louis, MI). D-Lin-MC3-DMA (MC3) was purchased from MedChemExpress (NJ, USA).

### siRNAs

Target sequences for siRNAs are listed in Table 1. All *in vitro* siRNAs were synthesized by Integrated DNA Technologies (IDT) (Coralville, IA, USA). siMod-UTR3 was designed as previously done (7) and synthesized by IDT (Coralville, IA, USA) as an RNA duplex. HPLC purified and high quantity siRNAs for *in vivo* work, siUTR3, siHel2, siN367 were synthesized by the RNA/DNA Synthesis core at the City of Hope (Duarte, CA), whereas siP, siScrM7 and siGFP were either synthesized by GenePharma (Shanghi, China) or IDT (Coralville, IA, USA). siGLO Red was purchased from Dharmacon™ (Lafayette, CO).

**Table 1.**
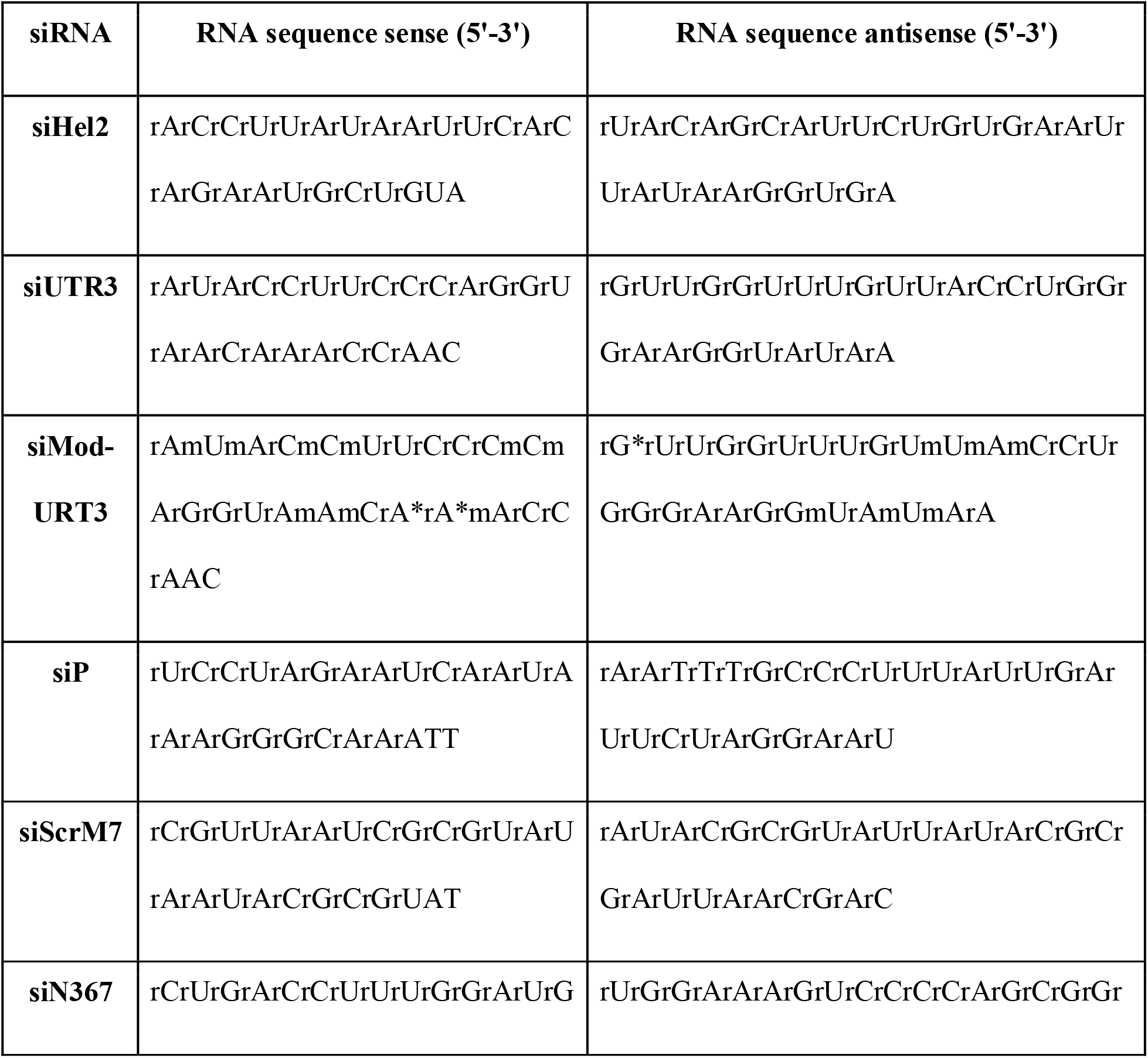
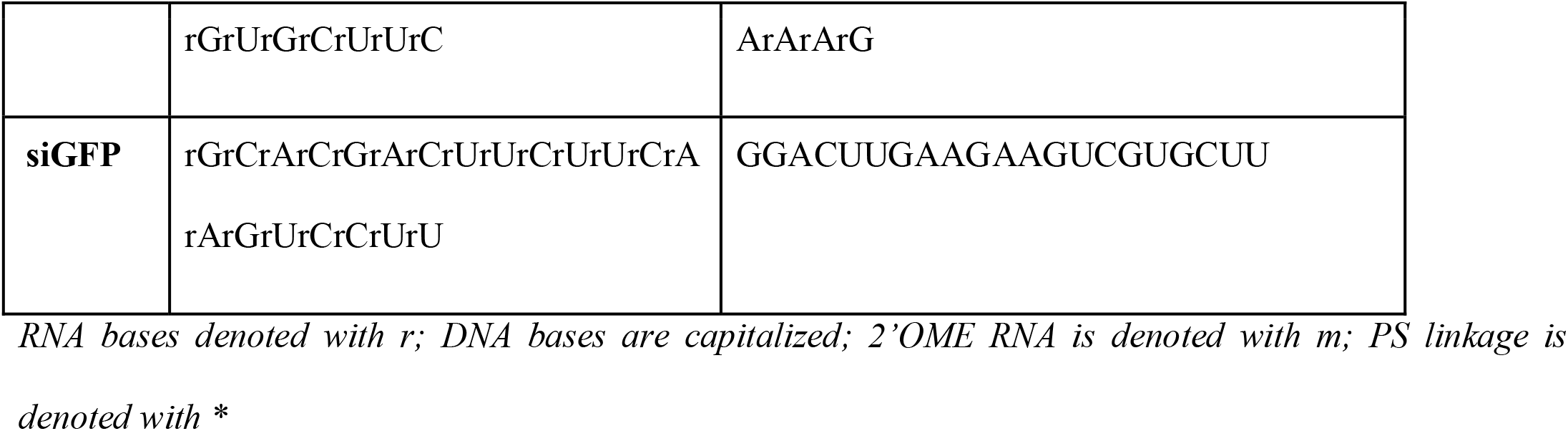
siRNA and D-siRNA sequences used in this study.

### Viruses

SARS-CoV-2 VOCs (Wuhan (Ancestral) – VIC1, B.1.351 (Beta) - VIC18383, B.1.617 (Kappa) - VIC18447, B.1.617.2 (Delta) - VIC18440, B.1.1.529 (Omicron) (BA.2 sub variant) - VIC35864) were obtained from the Peter Doherty Institute for Infection and Immunity and Melbourne Health, Victoria, Australia and cultured in Vero E6 cells. Viral supernatant was concentrated in Amicon ® Ultra-15 Centrifugal Filter units (Merck, Germany) and viral titre determined by the viral immunoplaque assay. RSV strain A2 (RSV-A2) was obtained from Professor Paul Young (School of Chemistry and Molecular Biosciences, The University of Queensland, QLD, Australia). RSV-A2 was cultured in A549 cells, ultracentrifuged, and purified on a sucrose gradient before determining viral titre using a viral plaque assay. *Viral plaque and immunoplaque assay*

For SARS-CoV-2 viral plaque assays, Vero E6 cells were infected with a MOI 0.002 (250 plaque forming units (PFU)) of SARS-CoV-2 for 1hr before overlaying with 1% methylcellulose- viscosity (4,000 centipoises) (Sigma- Aldrich, St. Louis, MO). Cells were incubated for 4 days at 37°C before fixing in 8% formaldehyde and stained with 1% crystal violet to visualize plaques. Viral immunoplaque assays for SARS-CoV-2 were performed on Vero E6 cells as described previously (11) using recombinant monoclonal antibodies that recognize SARS-CoV-2 (CR3022). Antibodies were obtained from Dr Naphak Modhiran and Associate Professor Dan Watterson (School of Chemistry and Molecular Biosciences, The University of Queensland, QLD, Australia). Viral immunoplaque assays for RSV-A2 were performed on A549 cells using anti-RSV-F rabbit polyclonal sera. Anti-RSV sera was obtained from Professor Paul Young (School of Chemistry and Molecular Biosciences, The University of Queensland, QLD, Australia). Virus titers are denoted as plaque forming units (PFU)/grams (g) of tissue.

### sLNP synthesis

Nucleic acid-entrapped PEGylated lipid particles were prepared as previously described (12). Briefly, required amounts of DOTAP, cholesterol, DOPE, and PEG2000-C16 Ceramide were dissolved individually in tert-butanol before being mixed at a molar ratio of 50:35:5:10. The required amounts of siRNA were dissolved in sucrose containing water (sucrose amount was calculated to generate an isotonic solution at the final step). A nitrogen: phosphate (N/P) ratio of four was generated by mixing the solutions mentioned above (1:1 v/v) to formulate the co-solvent system before the mixture was snap-frozen and freeze-dried (Alpha 1-2 LDplus, Martin Christ, Germany) at 0.05mbar overnight. The freeze-dried matrix was then hydrated with sterile nuclease free water to the required concentration. Nanoparticles were then filtered through a 0.45μm filter and stored at 4°C. The mixture was left at room temperature for 2h prior to characterization.

### dmLNP synthesis

Lipids were prepared at a 40:25:10:22:3 (DOTAP:MC3:DSPC:Chol:PEG) molar ratio. Lipids in ethanol were mixed with siRNAs in an aqueous phase at a mol cationic lipid: mol RNA (N:P) ratio of 3:1 using the NanoAssemblr Benchtop machine (Precision NanoSystems; Vancouver, BC, Canada). This machine contains a microfluidic chip by which the injected lipids and nucleic acids are mixed rapidly in a staggered herringbone pattern at a total flow rate of 12mL/min. The controlled mixing of the aqueous and organic streams produces homogeneous nanoparticles. Immediately following the mixing process, the nanoparticles were diluted 1:4 with 1xPBS to reduce the amount of ethanol present in solution. The nanoparticle solution was further diluted with 1xPBS up to 15mL and then concentrated in a 100kDa Amicon ultra-15 filter (Millipore; Burlington, MA, USA) via centrifugation at 400g for 120min. The flow through was discarded and another 15 mL 1xPBS was added to the column and centrifuged at 400g for 180min. Concentrated nanoparticles were then filtered through a 0.22μm filter and stored at 4°C.

### LNP characterization

The particle size and polydispersity index of the formulations were obtained using Zetasizer Nano ZS (Malvern Instruments, Malvern, UK) following appropriate dilution in PBS. All measurements were carried out at room temperature.

### In vivo infection and siRNA administration

All animal experiments were performed in compliance with relevant laws and institutional guidelines and in accordance with the ethical standards of the Declaration of Helsinki. For SARS-CoV-2 work, K18-hACE2 mice were purchased from the Jackson Laboratory (Bar Harbor, ME) and bred inhouse at the Griffith University Animal Resource Center. Mice were intranasally (IN) infected with 10^4^-10^5^ plaque forming unit (PFU) (20μL total volume) of live SARS-CoV-2 ancestral strain or delta VOC while under isoflurane anesthesia. Mice were subsequently treated with either naked, sLNP or dmLNP complexed siRNAs retro-orbitally (IV) (100μL total volume) or IN (20μL total volume) while under isoflurane anesthesia. For siRNA dosing on the day of infection (0 days post-infection (dpi)), siRNA was administered 2h prior to infecting mice with SARS-CoV-2. Mice were monitored daily for weighing and clinical scoring. This work was conducted in a BSL3 approved animal facility at Griffith University (Animal ethics approval: MHIQ/14/21/AEC). For RSV work, BALB/c mice were purchased from the Animal Resource Center (Perth, Australia). Mice were IN infected with 10^6^ plaque forming unit (PFU) (20μL total volume) of live RSV-A2 while under ketamine/xylazine anesthesia. Mice were subsequently treated with either sLNP complexed siRNAs via tail vein injection (IV) (200μL total volume) or IN (20μL total volume) while under ketamine/xylazine anesthesia. Cyclophosphamide was administered to mice intraperitoneally (IP) at a single dose of 100mg/kg five days prior to RSV-A2 infection as previously reported (13). Mice were monitored daily for weighing and clinical scoring. This work was approved by the Griffith University Animal Ethics Committee (Animal ethics approval: MHIQ/02/14/AEC).

### In vivo biodistribution studies

1,1□-Dioctadecyl-3,3,3□,3□-tetramethylindotricarbocyanine iodide (DiR) dye was used at a DiR: lipid ratio of 1 in 5400 (w/w) to track the lipid nanoparticles. siGLO Red (Cy3-labelled) transfection indicator (siGlo) was used as an indicator to track the LNP cargo (i.e., siRNA). The biodistribution of DiR-labelled LNP formulations and siGlo encapsulated LNPs were examined in extracted mouse organs on the PhotonIMAGER Optima *in vivo* imager (Biospace Lab, France) at the following wavelengths: excitation 720nm/ emission 790nm for imaging DiR, and excitation 547nm/ emission 563nm for imaging siGLO Red.

### Statistical analysis

All statistical analyses were performed using the statistical software package GraphPad Prism 9 and described in detail in respective figure legends.

## Results and Discussion

We have previously demonstrated highly potent *in vitro* and *in vivo* targeting of SARS-CoV-2 Wuhan (ancestral) strain with siRNAs targeting the ultra-conserved regions in the helicase (siHel2), and untranslated region (siUTR3) (7). Here, we tested these siRNAs against all SARS-CoV-2 variants of concern (VOC) and find that they are equipotent *in vitro* (Figure 1A). Next, we wanted to determine whether the potent antiviral effect observed previously with these siRNAs *in vivo* can be recapitulated in mice infected with another SARS-CoV-2 VOC. Given that siHel2 exhibited a lower IC_50_ than siUTR3 against the B.1.617.2 (delta) variant (Figure 1B), we complexed siHel2 with stealth LNPs (sLNPs) for IV administration, parallel to what we did previously for targeting SARS-CoV-2 ancestral strain *in vivo* (7). LNP systems are typically composed of a zwitterionic lipid, an ionizable cationic lipid, poly(ethylene glycol)-lipid (PEG-lipid), cholesterol, and the nucleic acid cargo. Unlike standard liposomes, we have developed sLNPs that are specifically formulated to be stable in serum, circulate for long periods, and protect siRNA payloads from nucleases (12, 14). Consistent with our previous observations (7), prophylactic administration of siHel2-sLNP via IV administration results in almost complete amelioration of viral infection in mice lungs (Figure 1C).

**Figure 1.**
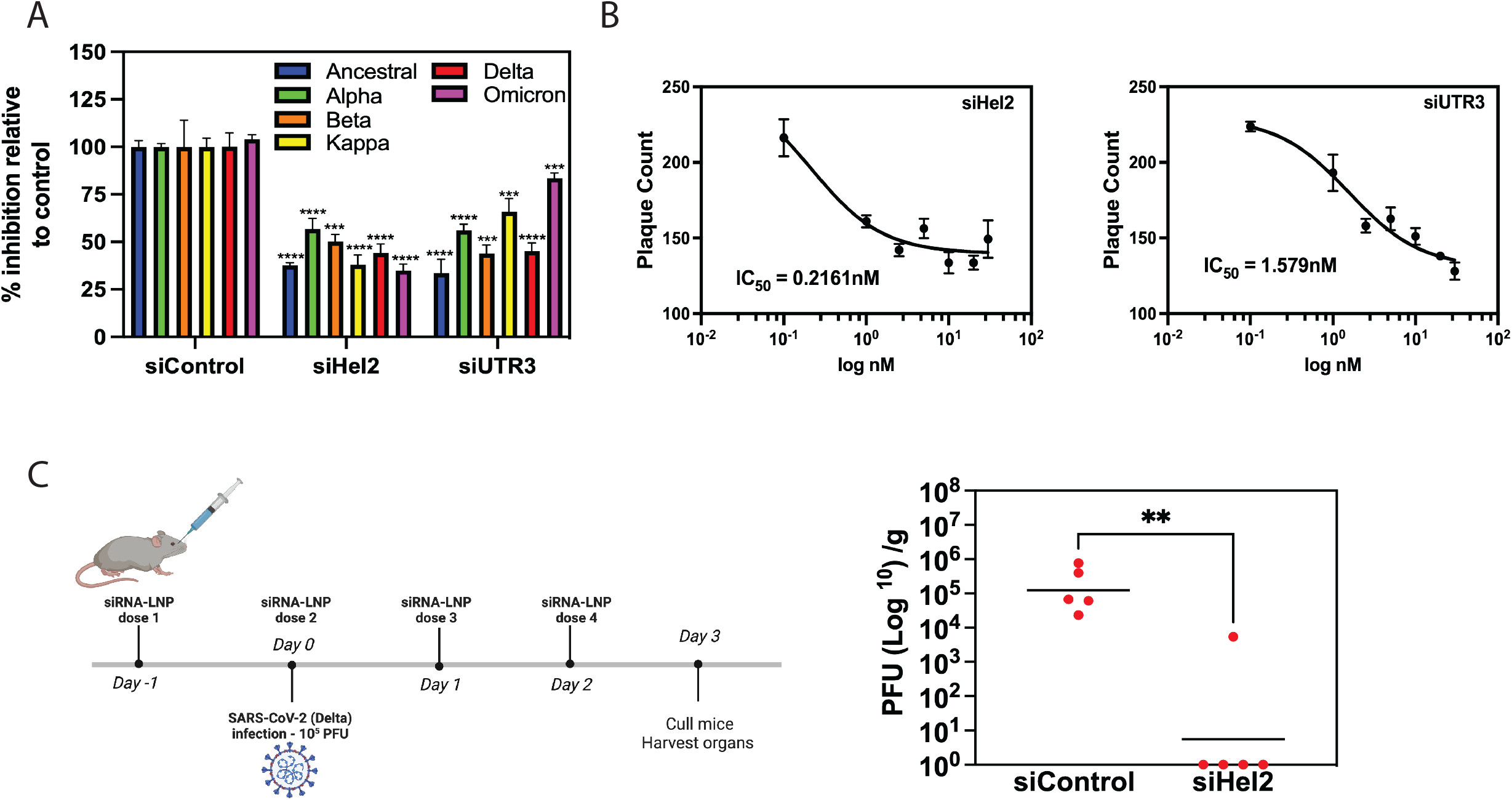
SARS-CoV-2 targeting siRNAs are effective at reducing lung viral infection *in vivo*. A) Vero E6 cells were transfected with control and targeting siRNAs (30nM) complexed to Lipofectamine 2000 for 24h before infecting with indicated SARS-CoV-2 VOC at 250 plaque forming unit (PFU). Infectious viral plaques were counted 4 days post infection (dpi). Data is expressed as mean percentage plaque inhibition relative to virus alone (control) and is representative of the standard error of the mean (SEM) of triplicate treatments. ∗p < 0.001 and ∗∗p < 0.0001, one-way ANOVA (Dunnett’s post-test) when compared against siControl. The control siRNA used here is siN367 (siControl). Data is representative of one out of three independent experiments. B) Vero E6 cells were transfected with increasing concentrations (0.1-30nM) of targeting siRNAs complexed to Lipofectamine 2000 for 24h before infecting with delta SARS-CoV-2 VOC at 250 plaque forming unit (PFU). Infectious viral plaques were counted 4dpi. Data is expressed mean plaque counts and standard error of the mean (SEM) of triplicate treatments and IC_50_ determined. C) K18h-ACE2 mice infected with 5×10^4^ live delta SARS-CoV-2 VOC received daily intravenous (IV) (retro-orbital) treatment of siRNA-sLNP (1mg/kg) daily (−1 to 2 dpi). The control siRNA used here is siN367 (siControl). Lung viral tissue counts/g at 3dpi are shown. Each dot represents data from one mouse and bars represent the mean. ** p<0.005 one-way ANOVA (Dunnett’s post-test).

Meyers *et al*., (15) previously screened a range of siRNAs targeting the phosphoprotein (P), N and large (L) protein genes and showed varying silencing potency against the RSV-A2 strain. Notably, the siRNA targeting the P gene (siP) exhibited over 95% inhibition at a concentration of 5nM. As was done previously with SARS-CoV-2 targeting siRNAs, siHel2 and siUTR3 (7), we converted RSV-targeting siRNA, siP, into dicer substrate-siRNA (D-siRNA) for more potent target mRNA silencing (Table 1). LNPs have been shown to functionally deliver nucleic acid payloads to the lung following IN administration (16). We postulate that IN administration of RSV-targeting siRNAs complexed in sLNPs would functionally repress RSV infection in mouse lungs. Here, siP is complexed into sLNP (siP-sLNP), as we have done previously for *in vivo* siRNA-specific SARS-CoV-2 lung targeting via the IV route (7). Indeed, prophylactic IV and IN administration of siP-sLNP resulted in significant reduction of lung viral load in RSV-A2 infected BALB/c mice at 3 days post-infection (dpi) (Figure 2A). Remarkably, we observed a similar effect with a therapeutic intervention regimen at 7dpi in mice pre-treated with cyclophosphamide (Figure 2B). Pre-treatment of BALB/c mice with cyclophosphamide, rendering the mice in an immunocompromised state, results in a more severe and sustained RSV *in vivo* infection model (13). Overall, both therapeutic and prophylactic IN and IV administration of RSV-targeting siRNA packaged into sLNPs results in the reduction of RSV lung infection *in vivo*. IN delivery of unmodified anti-RSV siRNA targeting the P gene have been attempted previously *in vivo* and resulted in effective reduction of RSV lung infection (17). Unmodified siRNAs are siRNAs that have not undergone necessary chemical modifications to protect it from nuclease degradation. It is important to note that the study administered almost double the amount of siRNA (70µg siRNA) we used here (40µg siRNA) and without the aid of a carrier system (e.g., LNPs). We speculate that this was to compensate for any loss of siRNA bioactivity due to their unmodified siRNAs becoming prone to nuclease degradation over time. Nonetheless, we show that targeting the RSV P gene prophylactically and therapeutically using targeting siRNAs encapsulated in sLNP can reduce RSV lung infection.

**Figure 2.**
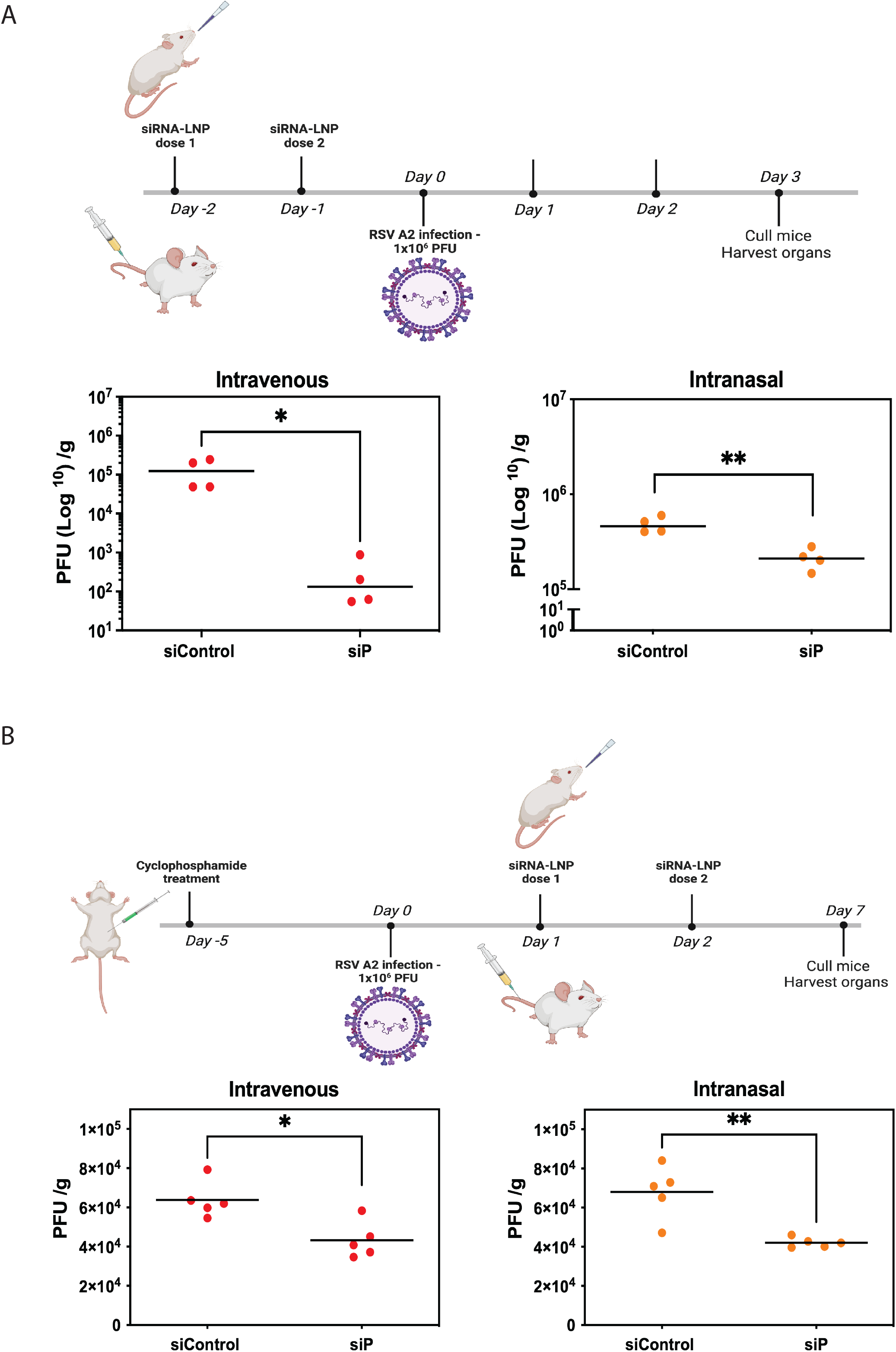
RSV targeting siRNAs are effective at reducing lung viral infection *in vivo*. A) BALB/c mice infected with 1×10^6^ live RSV-A2 received either IV (tail vein) or intranasal (IN) treatment of siRNA-sLNP (40µg siRNA) every 24h (−2 to -1dpi). The control siRNA used here is siScrM7 (siControl). Lung viral tissue counts/g at 3dpi are shown. Each dot represents data from one mouse and bars represent the mean. * p<0.05, ** p<0.005 one-way ANOVA (Dunnett’s post-test). B) BALB/c mice infected with 1×10^6^ live RSV-A2 received either IV (tail vein) or intranasal (IN) treatment of siRNA-sLNP (40µg siRNA) every 24h (1 to 2dpi). The control siRNA used here is siGFP (siControl). Cyclophosphamide was administered to mice intraperitoneally (IP) (100mg/kg) at -5dpi. Lung viral tissue counts/g at 7dpi are shown. Each dot represents data from one mouse and bars represent the mean. * p<0.05, ** p<0.005 one-way ANOVA (Dunnett’s post-test).

We have previously used LNP formulations composed of DOTAP with the ionizable lipid DLin-MC3-DMA (dmLNP) (18) and observed that both sLNPs and dmLNPs efficiently deliver siRNAs to the lungs of CoV-2-infected mice via IV administration (7). Interestingly, we observed that dmLNPs had better lung biodistribution than sLNPs when delivered via the IN route (Figure 3A) and that the majority of siRNA payload end up in the lungs compared to the nasal cavity when formulated with dmLNP (Figure 3B). Hence, we wanted to test this LNP formulation in an *in vivo* SARS-CoV-2 infection model by packaging siHel2 and siUTR3 in dmLNPs for IN delivery. Prophylactic IN administration of both SARS-CoV-2 targeting siRNA-dmLNPs reduced delta SARS-CoV-2 lung load but the effect was most significant with siUTR3 48h after the last siRNA dose (Figure 3C). We attribute this moderate effectiveness to the lack of siRNA biodistribution in mice the lungs at 48h post-IN delivery of siRNA-dmLNPs (Figure 3B). Indeed, this effect was demonstrably better when lung viral load was measured 24h after the last siUTR3 dose against both ancestral and delta SARS-CoV-2 variants (Supplementary Figure 1). Despite this, the observations suggest that strong anti-SARS-CoV-2 siRNA potency can be maintained *in vivo*, underscoring the effectiveness of siRNA-dmLNPs delivered via the IN route. Next, we then wanted to determine whether the observed anti-SARS-CoV-2 effect can be maintained when delivered without siRNA-dmLNP encapsulation. Notably, the previously observed antiviral effect was lost when siUTR3 was delivered unencapsulated (naked) by the IN route (Figure 3D). Though not significant, chemically modifying siUTR3 with 2’O-methyl and phosphorothioate modifications (siMod-UTR3), which have previously been shown to enhance siUTR3 stability and protection from nuclease degradation (7), maintained anti-SARS-CoV-2 effectiveness *in vivo* (Figure 3D). This data suggests that LNPs are important in protecting siRNAs and anti-SARS-CoV-2 activity *in vivo*. Indeed, Chang *et al*., (9) demonstrated lung anti-SARS-CoV-2 effect of IN-delivered chemically modified naked siRNAs *in vivo*, emphasizing the need to chemically modify siRNAs when delivering without a carrier vessel. Collectively, we find here that prophylactic IN administration of siRNA-dmLNPs can effectively reduce SARS-CoV-2 lung infection *in vivo*.

**Figure 3.**
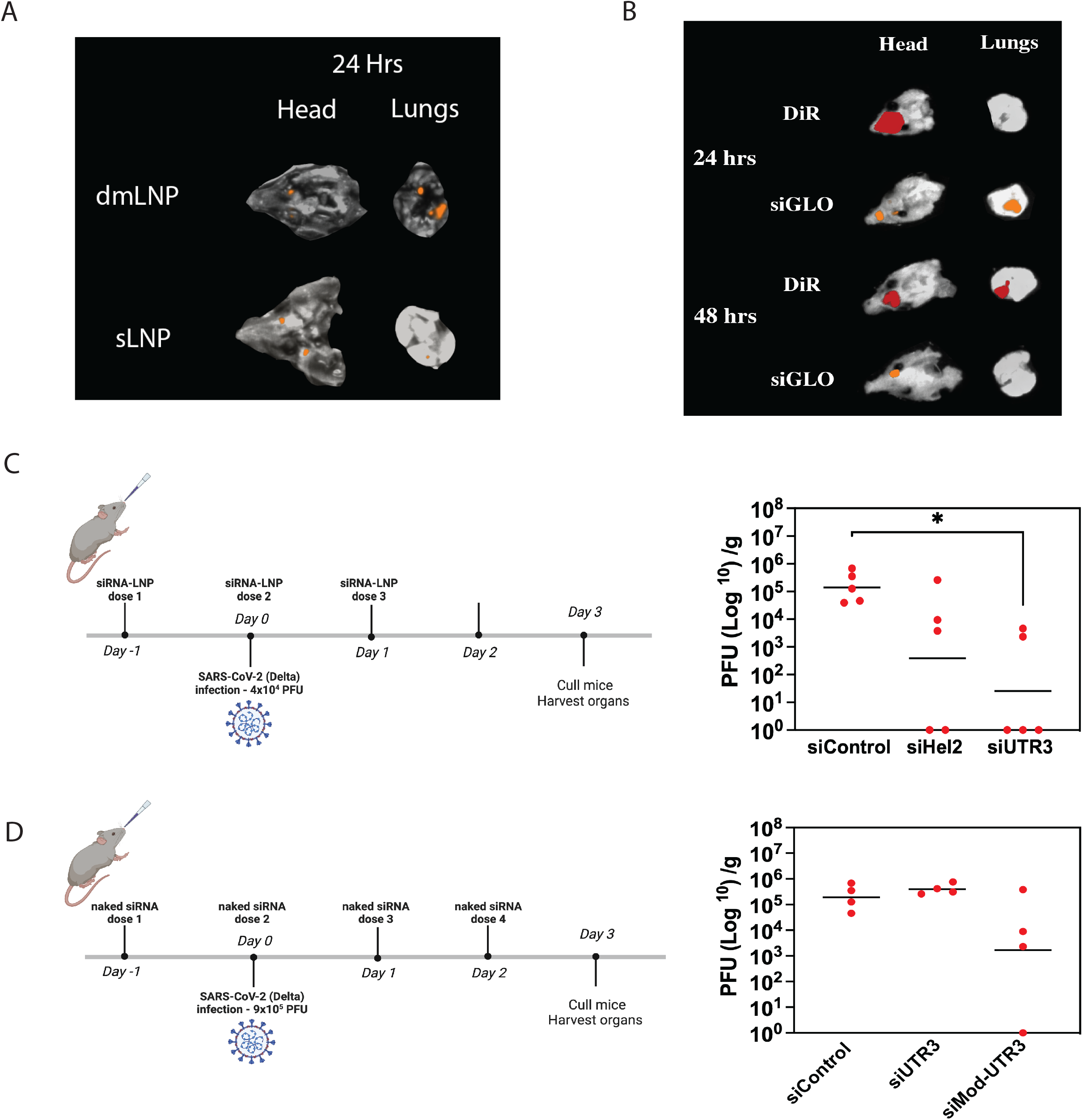
Intranasal delivery of SARS-CoV-2 targeting siRNAs reduces viral lung loads *in vivo*. A) K18hACE2 mice received either siGLO-dmLNP (1mg/kg) or siGLO-sLNP (1mg/kg) via IN treatment and organs harvested at various time points before measuring fluorescence on the PhotonIMAGER Optima *in vivo* imager. B) K18hACE2 mice received siGLO-dmLNP (1mg/kg) or dmLNP-loaded with fluorescent DiR via IN treatment and organs harvested at various time points before measuring fluorescence on the PhotonIMAGER Optima *in vivo* imager. C) K18h-ACE2 mice infected with 4×10^4^ live delta SARS-CoV-2 VOC received daily IN treatment of siRNA-dmLNP (1mg/kg) daily (−1 to 1dpi). The control siRNA used here is siN367 (siControl). Lung viral tissue counts/g at 3dpi are shown. Each dot represents data from one mouse and bars represent the mean. * p<0.05 one-way ANOVA (Dunnett’s post-test). D) K18h-ACE2 mice infected with 9×10^5^ live delta SARS-CoV-2 VOC received daily IN treatment of unencapsulated (naked) siUTR3, siMod-UTR3 (1mg/kg) or siControl-dmLNP (1mg/kg) daily (−1 to 2dpi). The control siRNA used here is siN367 (siControl). Lung viral tissue counts/g at 3dpi are shown. Each dot represents data from one mouse and bars represent the mean.

The impact of emerging respiratory virus infections in humans is not trivial. Indeed, there is currently a need for the development of antiviral RNA therapeutics to control virus infections especially those with pandemic capacity. A modular therapeutic platform that can treat respiratory RNA viruses, as well as their emerging variants could not only be paradigm shifting in the management of pandemics, but also in the preparedness for future pandemics. Additionally, there is an urgent need for new platforms that enhance RNA therapeutic efficacy, bioavailability and biodistribution. Importantly, siRNA delivery into the lower respiratory tract via the IN route poses several challenges (e.g., nasal mucosal barriers). While IV works to deliver siRNA to the lungs (7), it is not a practical approach nor amendable in a clinical outpatient setting. A more practical approach to deliver siRNA is by direct inhalation and/or nebulization. Here, we utilize LNPs as effective carrier vessels for siRNAs delivered via the IN route and showed that SARS-CoV-2 and RSV-targeting siRNAs were efficacious in infected mouse lungs. siRNAs are highly specific to their targets and are now in the clinic for several human disorders (19). For example, siRNA therapeutics developed by Alnylam Pharmaceuticals, patisiran (IV administered) and givosiran (subcutaneously administered), were both approved by the FDA in recent years to treat polyneuropathy. However, we are yet to see an approved inhalable siRNA therapeutic product, possibly due to the inherent challenges associated with this mode of delivery (i.e., effective biodistribution and mechanical method for administering an inhaled drug) (20). Though we show that IN delivered LNP encapsulated siRNAs are effective in respiratory virus infection models, whether these siRNA-LNPs can be aerosolized and maintain bioactivity in lungs thereafter has yet to be determined. Nonetheless, the data presented here sets precedence for delivering siRNA molecules using the LNP platform via the IN route for effective targeting of emerging respiratory viral infections in the lower respiratory tract.

## Acknowledgements

We would like to thank Dr Naphak Modhiran and Associate Professor Dan Watterson (School of Chemistry and Molecular Biosciences, The University of Queensland, QLD, Australia) for providing us with SARS-CoV-2 (CR3022) recombinant monoclonal antibodies. We would also like to thank Professor Paul Young (School of Chemistry and Molecular Biosciences, The University of Queensland, QLD, Australia) for providing us with RSV-A2 and anti-RSV sera and the Peter Doherty Institute for Infection and Immunity and Melbourne Health, Victoria, Australia for providing us with SARS-CoV-2 VOCs.

## Figure Legends

**Supplementary Figure 1.**
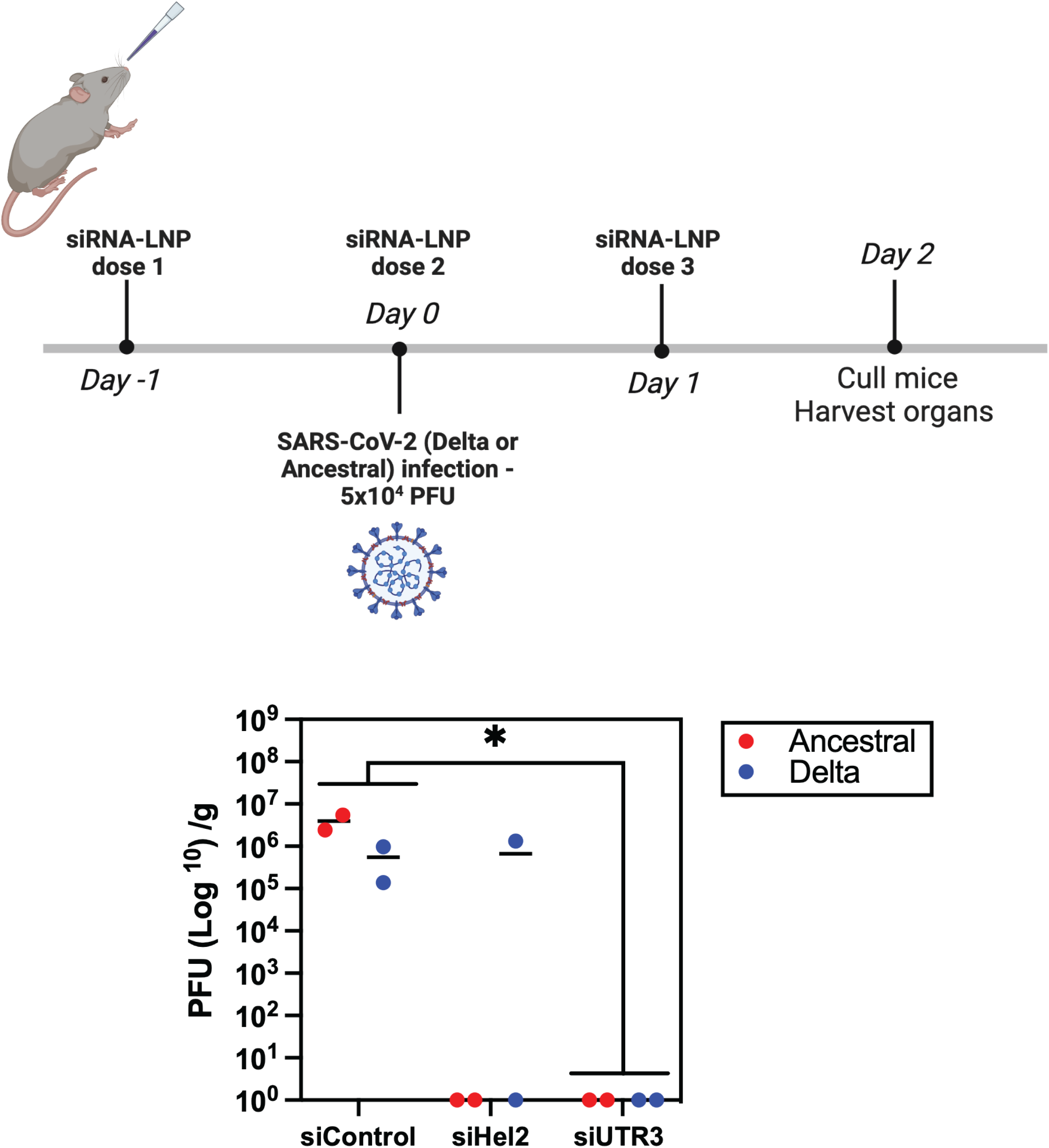
Intranasal delivery of SARS-CoV-2 targeting siRNAs reduces SARS-CoV-2 ancestral and delta lung loads *in vivo*. K18h-ACE2 mice infected with 5×10^4^ live ancestral, or delta SARS-CoV-2 VOC received daily IN treatment of siRNA-dmLNP (1mg/kg) daily (−1 to 1dpi). The control siRNA used here is siN367 (siControl). Lung viral tissue counts/g at 2dpi are shown. Each dot represents data from one mouse and bars represent the mean. * p<0.05 Kruskal-Wallis test.

